# Comparison of genomic-enabled cross selection criteria for the improvement of inbred line breeding populations

**DOI:** 10.1101/2023.03.17.533166

**Authors:** Alice Danguy des Déserts, Nicolas Durand, Bertrand Servin, Ellen Goudemand-Dugué, Jean-Marc Alliot, Daniel Ruiz, Gilles Charmet, Jean-Michel Elsen, Sophie Bouchet

## Abstract

A crucial step in inbred plant breeding is the choice of mating design to derive high-performing inbred varieties while also maintaining a competitive breeding population to secure sufficient genetic gain in future generations. In practice, the mating design usually relies on crosses involving the best parental inbred lines to ensure high mean progeny performance. This excludes crosses involving lower performing but more complementary parents in terms of favorable alleles. We predicted crosses with putative outstanding progenies (high mean and high variance progeny distribution) using genomic prediction models to assess the value of top progeny. This study compared the benefits and drawbacks of seven genomic cross selection criteria (CSC) in terms of genetic gain for one trait and genetic diversity in the next generation. Six CSC were already published and we have proposed an improved CSC that can estimate the proportion of progeny above a threshold defined for the whole mating plan. We simulated mating designs optimized using different CSC and 835 elite parents from a real breeding program that were evaluated between 2000 and 2016. We applied constraints on parental contributions and genetic similarities between parents according to usual breeder practices. Our results showed that CSC based on progeny variance estimation increased the genetic value of superior progenies by up to 5% in the next generation compared to CSC based on the progeny mean estimation (i.e. parental genetic values) alone. It also increased the genetic gain (up to 4%) and/or maintained more genetic diversity at QTLs (up to 4% more genic variance when the marker effects were perfectly estimated).

## Introduction

Plant breeders have two main objectives—derive high-performing varieties at each cycle and improve the mean genetic value of their germplasm so as to be able to generate superior varieties in future generations. The mating design, i.e. the choice of the set of parental lines to cross and the progeny size per cross, is critical to ensure both short- and long-term genetic gain. However, the number of candidate crosses is putatively very high while the number of crosses and progenies that can be experimentally tested is often limited.

Breeders can decide on the mating design by ranking crosses according to *cross selection criteria* (CSC) that estimate their ability to produce superior progenies for a given trait of interest. The simplest way to rank crosses is based on the expected mean genetic value of the progeny that, in turn, can be estimated by the mean additive genetic value of the parental lines, or the so-called parental mean (PM) criterion (Jinks and Pooni 1976). However, this criterion does not use information on the genetic variance of progeny derived from a cross (e.g. the progeny variance) and thus does not differentiate, among crosses of similar PM, those with a higher potential to generate extreme (transgressive) progenies, i.e. superior to the best parent and likely to provide higher genetic gain. Several attempts have been made to predict the potential of a cross to produce high means but also extreme genetic variance in the progeny.

The progeny/gametic variance for inbreds/outbreds depends on the complementarity of favorable alleles between parents and their probability of recombining during meiosis (Zhong and Jannink 2007). Indeed, considering two QTLs, when alleles are in coupling phase (i.e. one parent carries the two beneficial alleles while the other carries deleterious ones), recombination decreases the progeny variance, while recombination increases this variance in repulsion phase. Regarding QTLs along the whole genome, progeny variance increases with the level of polymorphism between parents. In the past, genetic values were estimated via phenotypic observations (phenotypic selection [PS]). Phenotypic and then genotypic distances were assumed to reflect parental genetic complementary and were used to predict cross progeny variance (Souza and Sorrells 1991; Bohn et al. 1999; Utz et al. 2001; Hung et al. 2012).

More recently, genomic prediction (genomic selection [GS]) was developed to estimate genetic values from genotypes (genomic estimated breeding value [GEBV]). GS uses a training population (TP) which is phenotyped and genotyped to estimate the effects of segregating genomic variants (markers). Assuming that marker effects are additive, the GEBV of one individual is the sum of its allele effects at every marker. Compared to PS, GS can reduce the generation interval in crops via genotyping—rather than phenotyping—using rapid cycles based on simulations (Bernardo and Yu 2007, Bernardo 2009, Lorenzana and Bernardo 2009, Heffner et al. 2010, 2011a and b). Depending on the species and the quality of the TP used to build the prediction model, GS can also increase the prediction accuracy (Lorenzana and Bernardo 2009, Heffner et al. 2011 a and b).

Genomic predictions offer a promising alternative to estimate progeny variance using marker effects and recombination rate estimates. The progeny distribution can be estimated by simulating progeny *in silico* (stochastic simulation), whereby recombination events of parental genomes are placed along chromosome sequences according to a recombination map (Bernardo and Charcosset 2006, Mohammadi et al. 2015). Simulation has the advantage of taking progeny size into account when computing quantiles, but it is compute intensive. Alternatively, progeny distribution can be predicted using analytical formulas. To do so, the progeny breeding value distribution is assumed to be Gaussian, which is expected for traits controlled by a very high number of variants with small effects. The Gaussian distribution is centered on the expected progeny mean (progeny mean= Parental mean; PM), which can be estimated from the mean of additive parental genetic values using PS or GS. A formula to predict inbred progeny variance derived from a cross between two inbred lines was reported by Lehermeier et al. (2017) based on marker effect estimates using GS and their co-segregation in progeny derived from a genetic map. Formulas were also derived to estimate three- and four-way cross progeny variance (Allier et al. 2019a) and to predict gametic variance in an animal breeding context (Santos et al. 2019).

Several CSC using progeny distribution estimates have been put forward, with each having strengths and weaknesses. One strategy consists in estimating the genetic value of the best inbred progeny that could be derived from a cross. Daetwyler et al. (2015) defined the optimal haploid value (OHV) corresponding to the genetic value of the progeny of a cross that would cumulate the most desirable alleles or haplotypes of parents at each position. OHV is fast to implement and the selection of crosses based on this value has been shown to increase both the genetic values and genetic diversity of the superior fraction of progeny at the next generation, as compared to progeny derived from PM-based selection of crosses (Daetwyler et al. 2015; Lehermeier et al. 2017). Note that there is a very low probability of observing OHV in progeny as a high number of beneficial recombination events would be needed, while avoiding all disadvantageous ones. Considering that the progeny size is generally limited, another CSC named expected maximum haploid breeding value (EMBV) was suggested by Müller et al. (2018). EMBV predicts the value of a cross as the expected mean of the K top progenies among D allocated to the cross.

Another strategy is to predict the average genetic value of a superior fraction of the progeny of candidate crosses. Schnell and Utz (1975) suggested ranking crosses based on the expected mean of an upper fraction q of their progeny. This CSC was named the usefulness criterion (UC), with UC = PM + i*h*σ, where i is the selection intensity corresponding to the fraction q of selected progenies, h is the square root of heritability and σ is the progeny variance in our context. Note that when using UC in a GS context, h² (and thus h) is usually set at 1 for GEBV, but further research would be required to be sure that this assumption has no influence on the results. As an alternative to UC, Wellmann (2019) and Bijma et al. (2020) suggested computing the value of a cross as the probability of producing progeny superior to a given threshold. This threshold can be extrapolated from historical genetic gains observed in the breeding program (Wellmann 2019), or it can be estimated as corresponding to the usual per-generation selection rate among progeny (Bijma et al. 2020). It can also simply be the genetic value of the best parental line.

Several studies compared the efficiency of those CSC in the short-term selection response (one generation) (Zhong and Jannink 2007; Lehermeier et al. 2017; Yao et al. 2018; Bijma et al. 2020). The findings showed that CSC based on progeny variance estimation could actually increase the genetic gain, even if the parental genetic values and progeny variance were not accurately estimated. Zhong and Jannink (2007) and Bijma et al. (2020) showed that the relative benefits of CSC based on progeny variance estimation compared to PM depends on the ratio between the variance of progeny standard deviations - var(σ) - and the variance of progeny means - var(PM) - in the list of candidate crosses. When var(PM) among crosses is highly superior to var(σ), PM alone is enough to predict the rank of crosses.

According to the breeder’s equation, genetic gain is proportional to the genetic diversity and selection intensity (Falconer and Mackay 1966). In a closed breeding program, i.e. with no external genitors involved, the diversity decreases as the selection efficiency increases. A further objective of the mating design is thus to maintain sufficient genetic diversity to ensure long-term genetic gain. Breeders empirically avoid crossing the most related genitors (Wartha and Lorenz 2021) while ensuring that a sufficient number of parental lines will contribute to the next generation. Several more advanced methods have been designed to balance the expected genetic gain and expected genetic diversity at successive generations when selecting genitors and/or crosses, e.g. by constraining the average genetic similarity of all selected parents (Toro and Perez-Enciso 1990; Meuwissen 1997; Jannink et al. 2010; Akdemir et al. 2019; Allier et al. 2019a). In any case, the sought after balance between the expected genetic gain and expected genetic diversity is not trivial to define. It depends on whether the objective is to optimize short- or long-term genetic gain (e.g. in a breeding or pre-breeding program).

This study was carried out to compare the genetic values and genetic diversity of top inbred progenies derived from optimized mating designs obtained using different CSC. The parental population included 835 historical (2000-2016) lines from the INRAE-AO winter bread wheat breeding program. We tested several previously published CSC (PM, OHV, EMBV, UC) and adapted two new ones from the literature that had never been tested per-se. From Wellmann (2019), we adapted PROBA, which consisted of ranking crosses based on the expected proportion of progeny superior to the best breeding line of the breeding program. From Bijma et al. (2020), we defined the UC3 criterion maximizing the expected value of a superior fraction of the whole progeny of the mating design, without any approximation or hypothesis. We compared genetic gain and diversity levels in the selected progeny when the QTL effects and positions were supposedly known, and also when the marker effects were estimated using a GBLUP model with observed parental phenotypes. Diversity constraints on parental contributions, i.e. minimal and maximal number of parents, crosses and progenies were chosen according to typical breeding practices.

## Material and Methods

### Parental populations

The founder population included 835 F_8_-F_9_ winter-type bread wheat lines developed and phenotyped between 2000 and 2016 by breeders from the French National Research Institute for Agriculture, Food and Environment (INRAE) and its subsidiary breeding company Agri-Obtentions (AO) (Ben Sadoun et al. 2020). They were genotyped with 35K SNPs (Ben Sadoun et al. 2020) representative of the TaBW280K array (Rimbert et al. 2018). For this analysis, the markers were filtered according to the missing data rate (< 5%), heterozygosity rate (< 5%) and minor allele frequency (> 10%) yielding 16,429 SNPs. Missing genotypes were imputed using the Beagle v4.1 algorithm (Browning and Browning 2007; Browning and Browning 2016) implemented in the synbreed R-package (Wimmer et al. 2012). The genetic values for yield of these 835 lines were estimated using the GBLUP model.

### Different tested scenarios

Simulations were carried out to take three parameter levels into account:

(1) the degree of selection for the trait of interest in the parental population:
  (1a) **unselected population:** we considered that the parental population composed of 835 historical breeding lines from INRA/A0 had never been selected for the trait of interest. QTL positions and effects were randomly assigned.
  (1b) **selected population:** an ancestral population created as in (1a) was further crossed and selected via three *in-silico* cycles to produce the parental population. Note that the genetic architecture of this population was the same as the corresponding unselected parental population.
(2) the accuracy of marker effect estimates:
  (2a) **TRUE**: QTL effects and positions were supposedly known (or perfectly estimated).
  (2b) **ESTIMATED**: marker effects were estimated by GS using parental phenotypes and removing QTLs in genotypes (**Figure 1**)
(3) The constraints to maintain genetic diversity in the breeding material:
  (3a) **CONSTRAINTS** on parental contributions and genetic distance between parents (see below, constraints C1 to C6)
  (3b) **NO CONSTRAINTS** (only constraints C1 and C2 were applied to the total number of progenies and the minimum and maximum number of progenies per cross)

We simulated the eight scenarios that are summarized in **Figure 1**. Note that the corresponding CONSTRAINT/NO CONSTRAINT and TRUE/ESTIMATED scenarios were simulated with the same parental population and genetic architecture.

**Figure 1:**
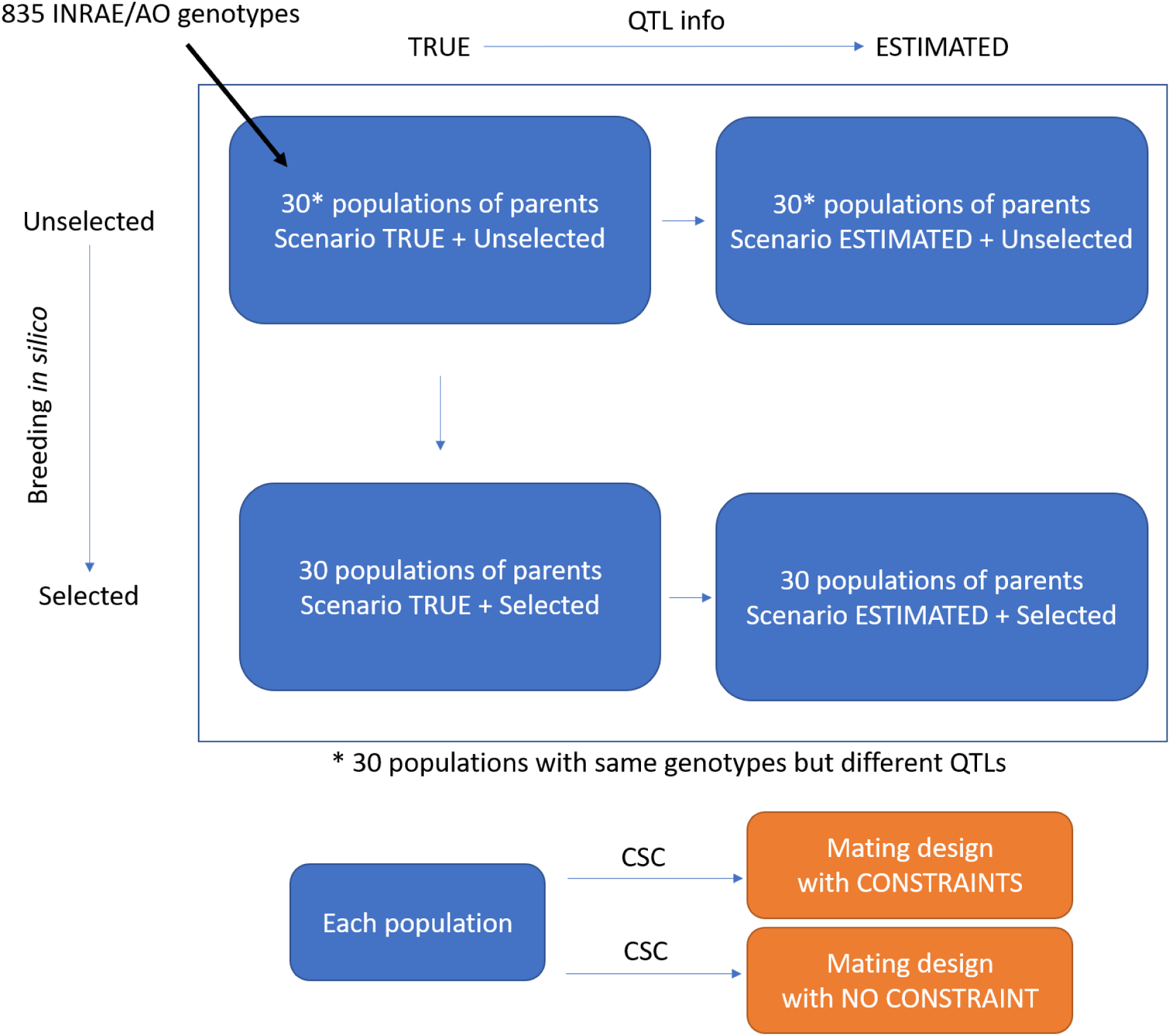
Different tested scenarios. The scenarios considered two marker effect estimation accuracy levels (TRUE, in which QTL effects were known and ESTIMATED, with marker effects being estimated by GS); two types of populations (unselected populations corresponding to the 835 INRAE/AO founders, and selected populations starting from those founders, followed by three random crossing and selection cycles); two mating design constraint levels (CONSTRAINTS and NO CONSTRAINTS). Each scenario was simulated for 30 different genetic architectures (characterized by a set of 300 QTLs with random position and effect) using INRAE/A0 historical breeding lines as the parental population.

#### Unselected population + TRUE QTL effects scenario

The parental population was built with genotypes of the 835 historical breeding lines from the INRAE/AO breeding program. In order to take into account the uncertainty in the genetic determinism of quantitative traits, we simulated 30 random genetic architectures controlled by 300 QTLs randomly picked among the 16k SNPs, with normally distributed genetic effects N(0,1). The favorable allele was assigned at random to one of the two SNP alleles, so that coupling and repulsion associations would also occur at random. QTL effects were adjusted to provide a variance of true breeding values (TBV) of 14 (quintal/ha)², as for the parental GEBV obtained using experimental data. TBV were calculated as the cross product between QTL effects and allelic doses.

#### Selected population + TRUE QTL effects scenario

Populations under selection for one or several traits of interest present negative covariances between QTLs. This phenomenon is called the Bulmer effect (Bulmer 1971). Hence, the observed genetic variance is lower compared to populations that have never been under selection. In unselected population simulations, this phenomenon was not taken into account as QTLs and effects were assigned at random positions along the genome. To take the Bulmer effect into account, we derived 30 ‘selected populations’ from the founders by applying three truncation selection cycles to the 30 unselected populations. At each of the three selection cycles, 300 crosses were performed at random from the 300 lines with the highest TBV. Selection on TBV provided an opportunity to maximize the Bulmer effect in new populations. Each cross produced 11 F5 RILs (total progeny = 3,300), simulated with the MOBPS R package (Pook et al. 2020). At cycles 1 and 2, only one progeny per cross was selected based on TBV. In the 3^rd^ cycle, the three best progenies per cross were kept, leading to a final population of 900 parental lines and called the ‘selected population’, from which 835 lines were sampled at random.

#### Unselected population + ESTIMATED QTL effects scenario

Phenotypes of unselected parents were simulated with a heritability ℎ²_0_ of 0.4 by adding a normally distributed noise of variance 21 (quintal/ha)² to their TBV (ℎ²_0_ = 14/(14+21) = 0.4).

Marker effects were estimated by backsolving the model using the PostGSf90 software package (Wang et al. 2012; Aguilar et al. 2014). GEBV of progenies were computed as the cross product between estimated marker effects and allelic doses.

#### Selected population + ESTIMATED QTL effects scenario

Phenotypes were simulated by adding a normally distributed noise of variance 21 (quintal/ha)² to the TBV. We used the same procedure as above to estimate marker effects and GEBV.

### Estimation of genetic values and marker effects

For the ESTIMATED scenarios, we used a GBLUP model to estimate parental line genetic values and marker effects according to the model:

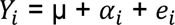

where i denotes the name of the parental line (n = 835), Y is the vector of phenotypes, μ is the average phenotype, α is the vector of genetic values and e is the vector of residual effects. The genetic values were assumed to follow 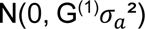, where G^(1)^ is the genomic relationship matrix computed as 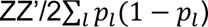, with Z being the centered genotyping matrix, excluding QTL genotype, and *p_l_* the allelic frequency at locus l, and where 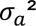 is the genetic variance. Residual effects were assumed to follow 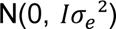. Parameters 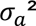 and 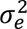 were estimated using the AIREMLf90 software package (Misztal 2008).

### Prediction of progeny variance

The expected variance of progeny was computed using the formula provided by Lehermeier et al. (2017) for biparental RIL progeny obtained after four generations of selfing (F5 RILs). For each cross *P_i_*\**P_j_*, the formula for the expected variance of progeny was

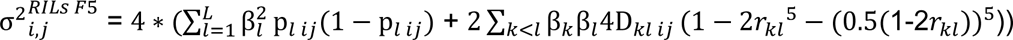

Where β are either QTL effects for TRUE scenarios (length β = 300) or estimated marker effects for ESTIMATED scenarios (length β = 16 429 - 300), p_*l ij*_ is the allelic frequency at locus l for parents *P_i_* and *P_j_* (0 if parents carry the same allele at this locus, 0.5 if they differ), D_*klij*_ is the linkage disequilibrium (LD) between alleles at loci l and k for parents *P_i_* and *P_j_* (either 0 if parents carry the same allele at locus l or k, or 0.25 if alleles are in coupling phase (i.e. one parent carries the two beneficial alleles while the other carries deleterious alleles), or – 0.25 if the alleles are in repulsion phase and *r_kl_* is the recombination rate between locus l and k. The recombination rates were computed from the Western European recombination map published by Danguy des Déserts et al. (2021), using the Haldane mapping function (Haldane and Waddington 1931): *r_kl_* = 0.5*(1 − *e*^−2*dkl*^), where *d_kl_* is the genetic distance (in morgans [M]) between loci k and l (Haldane, 1919).

The estimation of progeny variance for a high number of crosses (348,195 crosses in our study) and of simulations (n=120) was highly time consuming. We accelerated this estimation as described in **Supplementary Protocol S1**.

### Mating design constraints

Selecting crosses with the best CSC while including constraints on the progeny allocation across parents can be defined as an optimization problem in which variables to adjust (progeny sizes of each candidate cross in our case) will determine the value of the objective function to maximize (the sum of products of CSC values by progeny sizes in our case) but are also subject to constraints (e.g. the number of progeny per cross and per parent could be limited). When the equation system is linear for the variables to adjust, linear programming may be used to find the set(s) of variables that maximize the objective function. Otherwise, for more complex problems, heuristic algorithms such as genetic algorithms (GA) may be used to obtain a good (but not necessarily the best) problem solution.

A mating design was defined by a vector giving the number of progenies *D_ij_* allocated to each candidate cross *P_i_*\**P_j_*. The constraints were inspired from the bread wheat breeding program of the private company Florimond Desprez (personal communication):

- C1: The total number of progenies was set at D = 3 300
- C2: The number of progenies allocated to a cross ranged from *D_min_*= 5 to *D_max_* = 60
- C3: The number of crosses ranged from *K_min_*= 200 to *K_max_*= 300
- C4: The number of progenies derived from one parent could not exceed *C_max_* = 250
- C5: The number of recruited parents for the mating design ranged from *P_min_*= 100 to *P_max_* = 132
- C6: Highly related parental lines could not be crossed. We used the LDAK software package (Speed et al. 2012) to obtain a genomic relationship matrix G^(2)^ in which SNPs were weighted according to local linkage disequilibrium (LD) in order to take into account the very heterogenous LD in bread wheat, which markedly increases from telomeres to centromeres. This variance-covariance matrix was computed as WW’, where W was obtained by centering and scaling each column of the genotyping matrix Z such that *W_l_*= *w_l_* 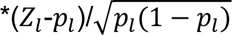 where *P_i_* is the allelic frequency at locus l and *w_l_* is the weight estimated by LDAK according to the local LD intensity. Crosses involving a pair of parental lines showing covariance superior to the 99% quantile covariance value were removed from the list of candidate crosses (1% of the candidate crosses).

We compared scenarios with and without constraints, i.e. respectively called ‘CONSTRAINTS’ and ‘NO CONSTRAINT’. Only constraints C1 and C2 were considered for the NO CONSTRAINT scenarios. Note that parental lines GEBV and estimates of marker effects were the same for the CONSTRAINTS and NO CONSTRAINT scenarios.

In summary, we compared the benefits of CSC for eight scenarios: two scenarios that differentiated the type of parental population (unselected or selected), two scenarios with different genomic prediction accuracies (TRUE or ESTIMATED) and two scenarios with different diversity constraints applied on the mating designs (CONSTRAINTS and NO CONSTRAINT).

### CSC and their corresponding objective function

One mating design is defined by a set of crosses and their respective number of progenies. For each CSC, the mating design maximizes a specific objective function under constraints C1 to C6 for the CONSTRAINTS scenarios and C1 to C2 for the NO CONSTRAINT scenarios.

#### PM (parental mean)

The usefulness of the *P_i_*\**P_j_* cross iss the expected progeny mean, estimated as

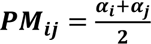

Where α is either the TBV of parents for TRUE scenarios or GEBV for ESTIMATED scenarios. The objective function to maximize is

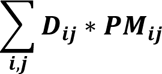

#### UC1 (usefulness criterion 1) (Schnell and Utz 1975)

This CSC is the expected mean of the q = 7% best progeny of a cross, computed as

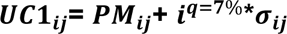

Where *i^q=7%^* ∼ 1.91 is the selection intensity corresponding to a 7% selection rate (computed as the inverse Mills ratio) and σ_*ij*_ is the progeny standard deviation. The progeny standard deviation σ_*ij*_ is computed either with QTL effects for TRUE scenarios or estimated allelic effects for ESTIMATED scenarios. Note that a 7% selection rate is usually applied at the Florimond Desprez company between F5 and F6 generation (when genomic predictions are applied). The objective function to maximize is

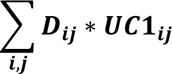

#### <u>UC2 (</u>usefulness criterion 2)

This CSC is the expected mean of the q = 0.01% best progeny of a cross, computed as:

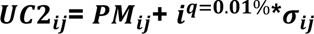

Where *i^q=0.01%^* ∼ 4 is the selection intensity corresponding to a 0.01% selection rate, i.e. twice the selection intensity of the UC1 criterion. Although this 0.01% selection rate is not realistic considering the small progeny size (*D_max_* = 60 progenies per cross), the objective is to select crosses with higher expected genetic variance compared to the UC1 criterion, while counting on them providing more outstanding progenies. The corresponding objective function to maximize is:

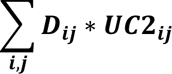

#### EMBV (expected maximum breeding value) (Müller et al. 2018)

The expected value of the best progeny among *D_ij_* allocated to a cross is:

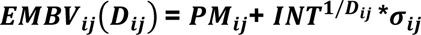

With *INT*^1/*Dij*^ being the expected value of the highest order statistic among a sample of *D_ij_* statistics drawn from N(0,1). An approximation of *INT*^1/*Dij*^ was provided by (Burrows 1972):

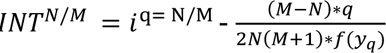

Where f is the density function of a Gaussian law N(0,1) and *y_q_* is the truncation threshold such that P(*y* ≥ *y_q_*) = q = N/M. In our conditions, N =1 and M = *D_ij_* and *i*^q=N/M^ = *f*(*y_q_*)/*q*, so the formula of Burrows yields:

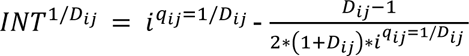

The objective function to maximize is:

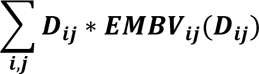

#### PROBA

This criterion ranks crosses based on their ability to produce a progeny exceeding a threshold λ, as suggested by Wellmann (2019) and Bijma et al. (2020). For setting λ we use the genetic value (TBV for TRUE scenarios or GEBV for ESTIMATED scenarios) of the best parental line. The probability of a *P_i_*\**P_j_* cross producing progeny with a genetic value superior to λ is 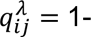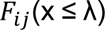, with *F_ij_* the cumulative distribution function of the Gaussian distribution 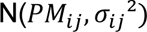.

The probability that no progeny of the *P* \**P* cross exceeds λ is 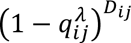. The probability that no progeny from all crosses exceed λ is 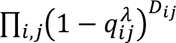, so the log probability is 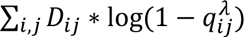. Maximizing the probability that at least one offspring will have genetic value greater than λ is equivalent to minimizing the objective function

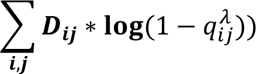

#### <u>UC3 (</u>usefulness criterion 3)

This criterion aims to maximize the expected mean of the superior quantile q (e.g. q = 7%) of progenies of the whole mating design, where q is the usual proportion of selected progenies. The same selection threshold *s_q_* is applied to all crosses and corresponds to the superior quantile q of the progeny genetic value distribution. The expected proportion of progeny of genetic value superior to *s_q_* differs for each cross and the total proportion of progeny exceeding *s_q_* is equal to q:

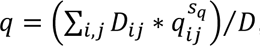, where 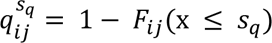 is the expected proportion of progeny superior to *s_q_* within the *P_i_*\**P_j_* family. The expected value of progeny superior to *s_q_* within each family is equal to 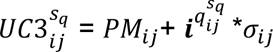. For a given mating design, as defined by the vector of *D_ij_*, the expected value of the q best progenies is thus equal to

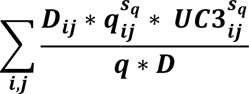

The best mating design is obtained by maximizing this objective function, with the constraint 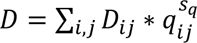.

#### OHV (optimal haploid value)

Daetwyler et al. (2015) defined OHV as the value of the best inbred progeny that could be theoretically derived from a cross. For each genomic segment b, the effects of haplotypes carried by parents *P_i_* and *P_j_* are respectively called β_*bi*_ and β_*bj*_. The OHV of a cross is computed as:

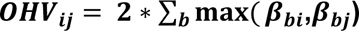

Daetwyler et al. (2015) showed that selecting crosses based on OHV instead of PM was advantageous in terms of short-term genetic gain when the number of haplotypic blocks per chromosome was low. For bread wheat, they showed, by simulation, that one to three blocks per chromosome allowed higher genetic gain than smaller blocks. We defined three haplotypic blocks per chromosome, one block per chromosome arm plus one block for the centromere (with the positions of centromeric regions defined in Choulet et al. 2014).

The objective function to maximize is:

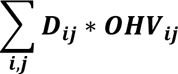

### Optimization of mating designs

In the CONSTRAINTS scenarios, for all CSC but EMBV and UC3, the objective function and constraints constituted a system of linear equations. We used an integer linear programming algorithm implemented in IBM ILOG CPLEX software (CPLEX Python API, IBM 2017) to maximize (or minimize) objective functions while respecting the constraints.

For criteria EMBV and UC3, the objective function and constraints did not form a system of linear equations, as the usefulness (e.g. the CSC value) of a cross actually depended on the number of progenies allocated to the cross. To optimize mating designs for EMBV and UC3 criteria, we used a genetic algorithm (GA). GAs are population-based metaheuristics inspired by Darwinism (Goldberg 1989). The GA description used in this study and the tuning parameters are given in **Supplementary Protocol S2**. GAs are difficult to tune and often remain stuck at local minima. To avoid premature convergence, a sharing process can be added before selection (Yin and Germay 1993) in order to give more chance to candidates that are isolated in the search space. The sharing process requires the definition of a distance between candidate solutions. Candidate solutions were considered different if at least one *D_ij_* was different. The population of candidate solutions per iteration was set at 100. At the first GA iteration, half of the initial candidate solutions were drawn at random, and the other half was set at linear programming optimization outputs of other CSC. The findings of a short preliminary study actually suggested that linear programming outputs of UC1 for EMBV optimization and PROBA outputs for UC3 optimization were the best starting points for EMBV and UC3 optimization.

For all criteria, we tested whether the pre-selection of candidate crosses with the highest PM would influence the value of the objective function to be maximized (**Supplementary Table S1**). For all criteria, pre-selection of the 10% highest PM crosses usually provided an objective function value after optimization that was 99% similar to the objective function value of the same population without pre-selection. To reduce the computation time, we thus optimized the mating designs with the 10% highest PM crosses. Note that pre-selection of crosses based on parental genetic values was also used in Zhong and Jannink (2007), Lehermeier et al. (2017) and Bijma et al. (2020).

In the NO CONSTRAINT scenarios, we did not use optimization software to optimize mating designs, except for UC3. For all other CSC, crosses were ranked based on CSC values and the 55 best crosses received *D_max_* = 60 progenies (constraint C2), for a total of D = 3,300 progeny (constraint C1).

### Progeny simulation

The F5 RIL progenies of each mating design were simulated using the MOBPS R package (Pook et al. 2020). Each mating design was simulated 20 times to account for the possibility that progeny genotypes might vary due to Mendelian gamete sampling. Progeny TBV were then computed as the cross product between QTL effects and the allelic dosage at QTL loci.

### CSC performance

The mating design optimization in this study had two objectives: to derive high-performing genotypes for commercial purposes, but also to improve the breeding population while limiting the loss of genetic diversity.

The ability of CSC to improve genetic values of commercial lines compared to PM at each K/D selection rate was computed as the relative increase in the mean progeny TBV compared to

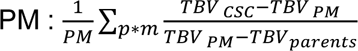

Where *TBV_CSC_*iss the mean TBV of the K best progenies among D simulated progenies in the m-^th^ simulation (M = 20 repetitions) of a mating design optimized for the CSC for the genetic architecture p (P = 30 different genetic architectures) for scenario s. The progeny selection rate (K/D) ranged from 1/3,300 (the very best progeny) to 10%. The term TBV stands for the mean TBV of all candidate parents in architecture p simulation m.

Finally, we measured genetic diversity using genic variance, computed as 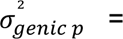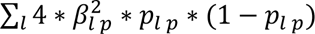, with *p_l p_* being the allelic frequency of QTLs in the selected progenies derived from population p in scenario s, and β_*l p*_ being the true allelic effect of QTLs at locus l in population p (note that QTL effects β did not change between scenarios, only between genetic architectures). The relative change in genic variance in the K/D selected progeny obtained using a mating design optimized for CSC compared to PM in scenario s was calculated as the relative increase in the progeny variance compared to PM:

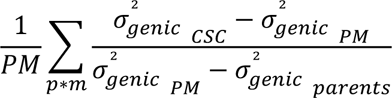

Where 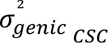 was the genic variance in the selected set of progenies in the m-^th^ progeny simulation for architecture p. To evaluate the ability of CSC to improve the new breeding population in terms of both gain and diversity, we set a 7% selection rate, corresponding to a realistic selection rate at the F5 stage in a bread wheat breeding program, and computed the relative increase in the mean progeny TBV and the relative increase in the progeny genic variance.

## Results

### Elite progenies

Crosses were selected using seven genomic cross selection criteria (CSC), namely PM (parental mean value = expected progeny mean value), UC1 (expected mean value of the top 7% progeny), UC2 (expected mean value of the top 0.01% progeny), UC3 (expected mean value of progeny superior to the 93% quantile of the whole mating design), PROBA (expected percentage of progeny superior to a threshold, set to the best parent value in this study), EMBV (expected value of the best progeny among D progenies) and OHV (best theoretical progeny value). They were computed with TRUE or ESTIMATED marker effects and using parents from unselected or selected populations for 30 different trait architectures. **Figure 2** gives the relative increase in the mean progeny TBV compared to the PM reference criterion for a selection rate ranging from 0.03% (selection of the best progeny among D = 3,300 progenies) to 10%.

**Figure 2:**
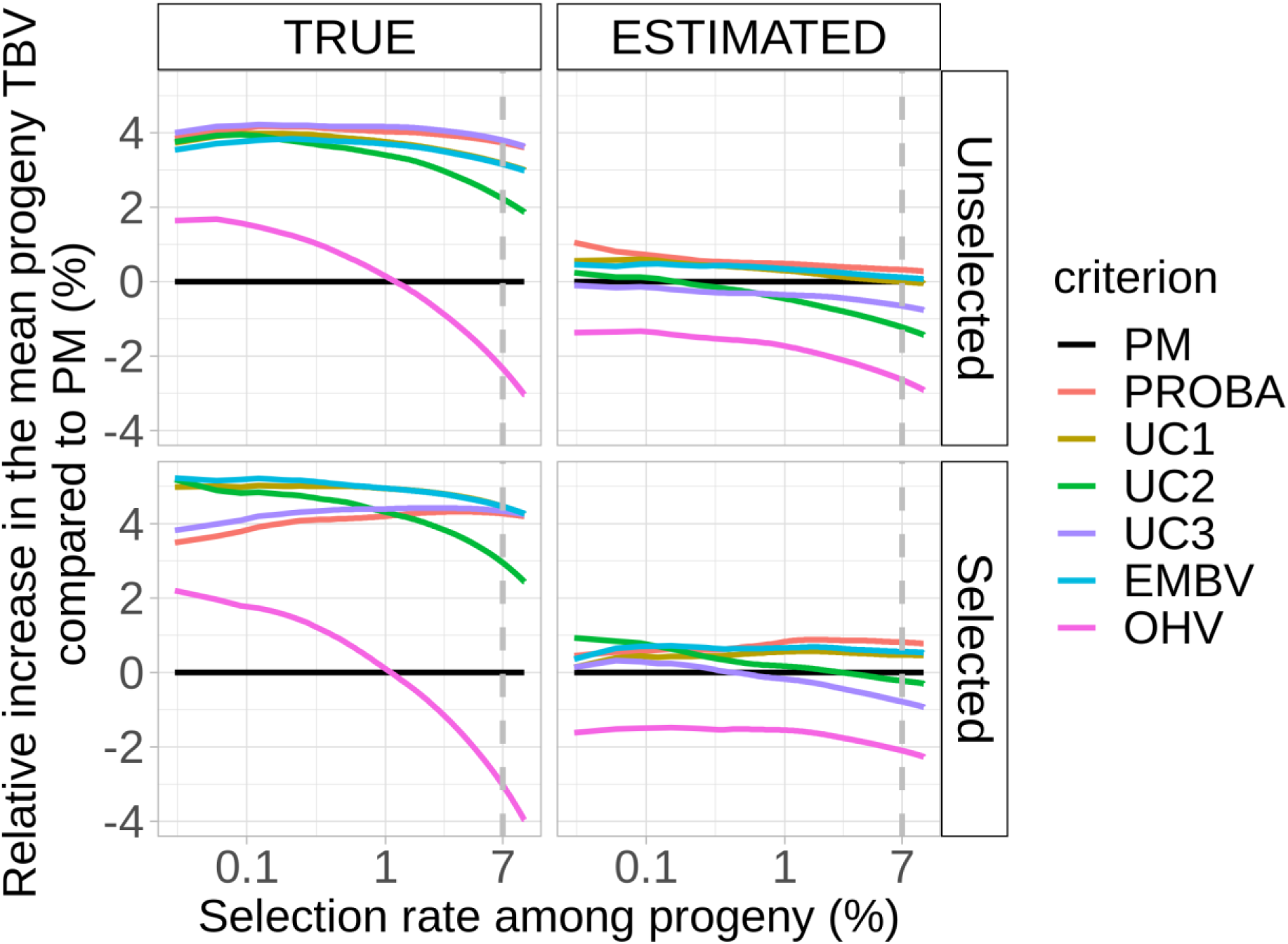
Relative increase (%) in the mean TBV of selected progeny obtained by optimizing the mating design with alternative CSC compared to the PM criterion. The vertical grey dashed line represents a 7% selection rate, as used in Figure 3.

For TRUE scenarios, all criteria were superior to PM when selection was strong (selection rate < 1%). For example, for all CSC but OHV, the mean TBV of the selected progeny increased by around 4% for unselected scenarios and by up to 5% for selected scenarios compared to PM. For ESTIMATED scenarios, the relative increase barely exceeded 1% for all scenarios.

The ranking of criteria to maximize the TBV of selected progeny changed slightly with the scenario and selection rate. For TRUE + unselected scenarios, the best criterion to maximize the value of the best progeny was UC3, with a 4.1% average increase in the TBV of the best progeny and a 1.9% standard deviation; for TRUE + selected scenarios, the best criterion was EMBV (5.2% ± 1.7%); for ESTIMATED + unselected scenarios, the best criterion was PROBA (1.1% ± 3.1%); and for ESTIMATED + selected scenarios, the best criterion was UC2 (0.9% ± 2.9%). Pairwise t-tests computed within each of the four scenarios identified three significant groups (p < 5% after Bonferroni correction) for TRUE scenarios: the upper group consisted of UC1, UC2, UC3, PROBA an EMBV, and the middle group consisted of PM and the lower group consisted of OHV. The pairwise t-tests were not significant for the ESTIMATED scenarios, except for OHV, which was significantly lower than the other CSC. In conclusion, CSC alternatives to PM (except OHV) were superior to PM only for TRUE scenarios, with no substantial differences between them.

Note that PROBA and UC3 slightly underperformed for TRUE + selected scenarios when selection was strong (low selection rate). Other CSC such as UC1 or UC2 should be preferred in that case. For all scenarios, the OHV criterion provided the lowest genetic gain. It was very disadvantageous compared to PM for all scenarios at > 1% selection rate.

In conclusion, when QTL effects were perfectly estimated (TRUE scenarios), CSC based on progeny variance estimation (UC1, UC2, UC3, EMBV, PROBA) could increase the genetic gain by up to 5% in breeding programs.

### Trade-off between genetic gain and genetic diversity in selected progeny

We considered that the new breeding population included the 7% best progeny derived from the optimized mating design. **Figure 3** shows the relative increase in the mean progeny TBV and the genic variance in the new breeding populations using CSC instead of the PM criterion. The grey line in **Figure 3** shows criteria with the best trade-off between genetic gain and genic diversity. For all scenarios (TRUE/ESTIMATED; unselected/selected), PM was not amongst the best trade-offs. For example, for TRUE + selected scenarios (bottom left in **Figure 3**), crosses could be selected based on EMBV (blue point) or UC1 (yellow point), with these two CSC reducing the loss of genic variance within the 0 to 4% range compared to PM (black point). In fact, most criteria maintained more genic diversity than the PM criterion, except PROBA and UC3 for most scenarios.

**Figure 3:**
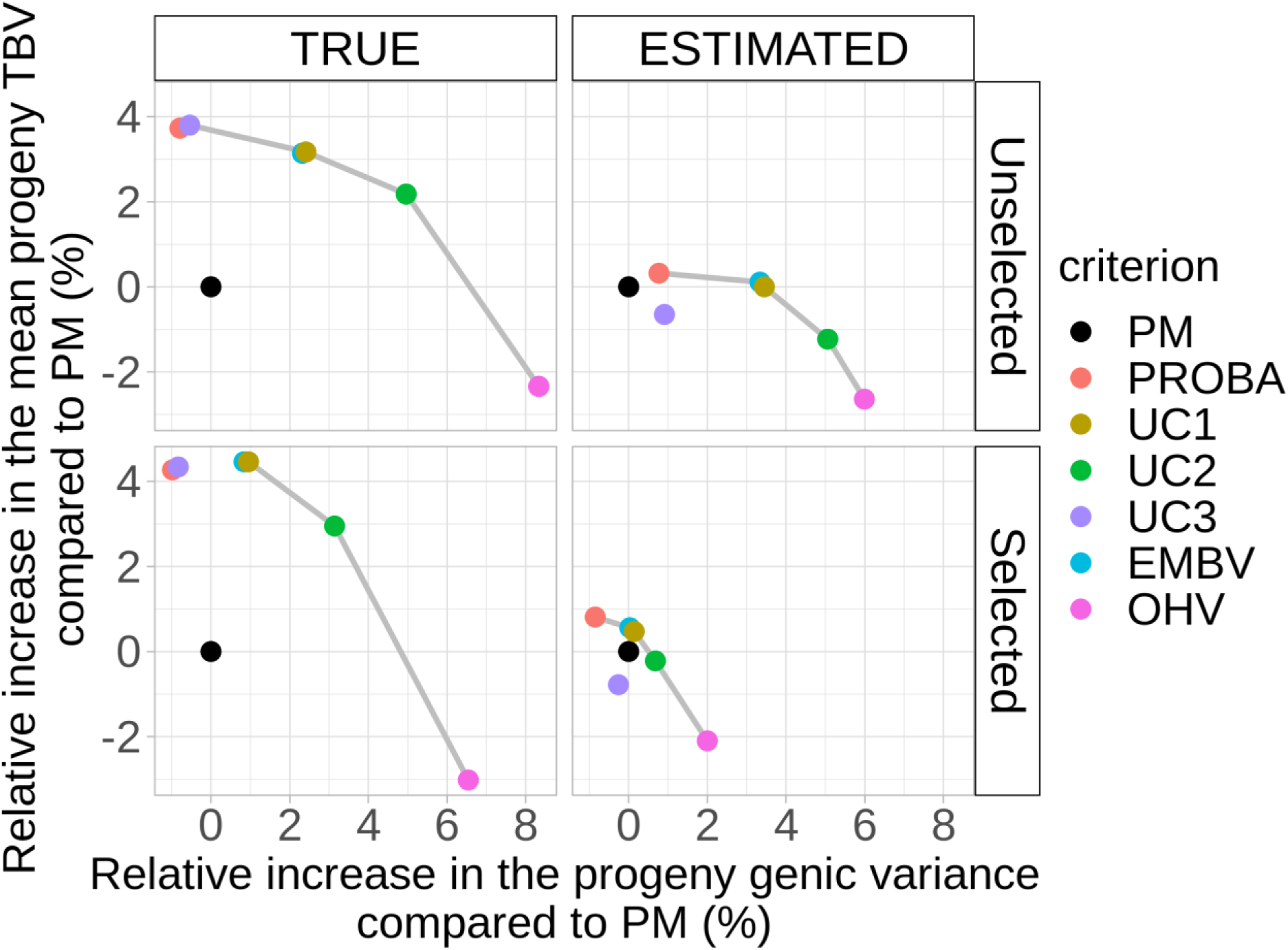
Trade-off between the relative increase in the mean progeny TBV and progeny genic variance in the 7% best progenies of the mating design compared to PM for CONSTRAINTS scenarios. Grey lines link criteria belonging to the set of best trade-offs, i.e. the best relative increase in the mean TBV for each level of relative increase in genic variance.

The set of criteria providing the best trade-offs was similar for all scenarios and included the OHV, UC2 and UC1 criteria, and sometimes PROBA. There was a negative relationship between genetic gain and genetic diversity. For example, OHV was the most efficient criterion to maintain genetic diversity but the worse one to maximize genetic gain, while PROBA was the opposite.

Genetic diversity within progeny depended on the diversity of the selected parents and the progeny distribution across the selected parents and crosses. Mating designs based on PM systematically displayed the highest average genetic similarities between selected parents compared to other CSC (**Supplementary Protocol S3, Supplementary Figure S1**). OHV and UC2 criteria displayed the lowest genetic similarities between recruited parents.

We used some CONSTRAINTS when building the mating design in order to maintain the genetic diversity, e.g. avoiding crossing related lines and ensuring that progeny were spread among a minimum number of parents and crosses. These CONSTRAINTS increased the genic variance by 10% (8-15% depending on the CSC and scenario) in the new breeding population and reduced the mean value of the new breeding population by 5% (4-8%; **Table 1**). Incidentally, the CONSTRAINTS also reduced the value of the top progeny by around 2% for all CSC and up to 8% when using PM.

**Table 1:** Impacts of CONSTRAINTS in terms of genic variance and genetic gain for the top 7% progeny of the whole mating design and for the best progeny value. The values were computed as 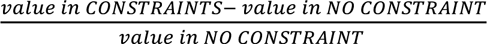 for each metric and each CSC, and then averaged over the 30 genetic architectures. Values in bold represent CSC showing the most desirable response in the CONSTRAINTS scenarios.

| Marker effects | TRUE |  | ESTIMATED |  |
| --- | --- | --- | --- | --- |
| Starting population | Unselected | Selected | Unselected | Selected |
| Genic variance of the breeding population | **PM + 12%**<br>Other CSC + 9% | **PM + 11%**<br>Other CSC + 8% | **PM + 15%**<br>Other CSC + 12% | PM + 13%<br>Other CSC + 13% |
| Genetic gain of the breeding population | PM: -8%<br>**Other CSC -6%** | PM: -8%<br>**Other CSC -5%** | **PM -4%**<br>Other CSC -5% | PM -4%<br>Other CSC -4% |
| Value of the best progeny | PM -8%<br>**Other CSC -2%** | PM -8%<br>**Other CSC -2%** | PM -4%<br>**Other CSC -2%** | **PM -1%**<br>Other CSC -2% |

The CONSTRAINTS scenarios had a significant effect for all three metrics (higher genic variance, lower genetic gain and lower best progeny value), especially when using PM. This could be explained by the fact that the selected crosses using CONSTRAINTS are sub-optimal compared to NO CONSTRAINTS in terms of genetic gain, while forcing a minimum level of diversity in parents. For NO CONSTRAINT scenarios, the algorithm assigns a maximum number of progenies (60) to the 55 best crosses, while for CONSTRAINTS scenarios, the objective function maximizes the sum of CSC of all selected crosses, with a limit of 250 progenies per parent for the whole design. The mate allocation concerning least performing parents seems more or less random with CONSTRAINTS (see the low percentage of crosses that were similar in two independent mating design optimizations from the same sets of parents using the PM criterion with CONSTRAINTS in **Supplementary Table S1**).

We observed an increase in the relative gain provided by CSC compared to PM in the CONSTRAINTS scenario. For example, in the TRUE + NO CONSTRAINT scenario, the relative increase in the TBV of top progenies was roughly twofold lower in comparison to the TRUE + CONSTRAINTS scenarios (**Figure 4**).

**Figure 4:**
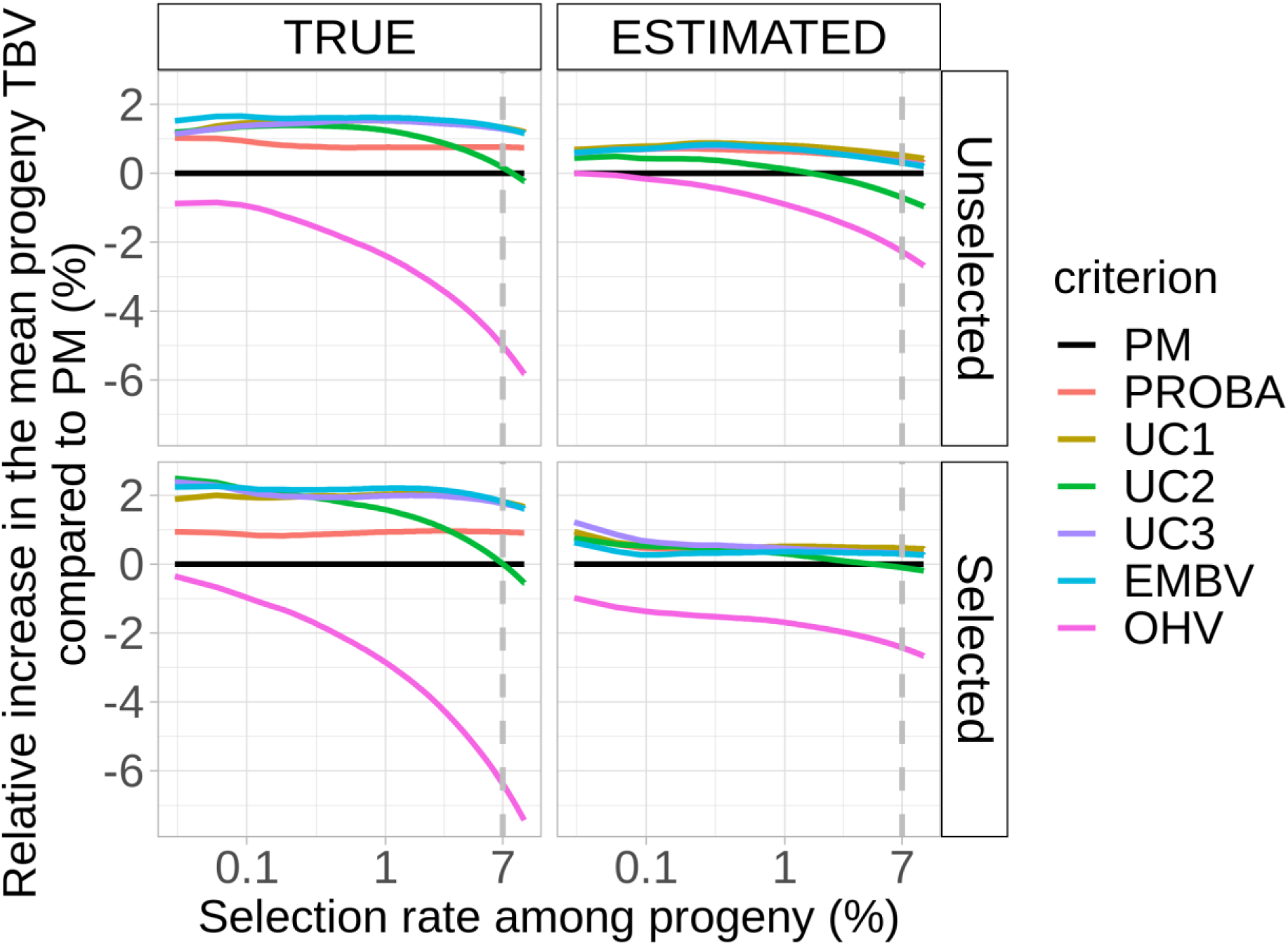
Relative increase in the mean TBV of selected progeny compared to progeny using the PM criterion for NO CONSTRAINT scenarios.

Despite the fact that the difference between PM and other CSC was reduced for NO CONSTRAINT scenarios, alternative CSC still appeared to be much more advantageous compared to PM in providing high-value progenies. For the TRUE + unselected scenario, EMBV provided the best progeny (relative increase compared to PM = 1.5% ± 1.6); for the TRUE + selected scenarios, UC2 provided the best progeny (2.9% ± 2.4); for the ESTIMATED + unselected scenarios, PROBA was the best criterion (0.7% ± 2.3) and for the ESTIMATED + selected scenarios (closer to breeding programs), UC3 was the best criterion (1.3% ± 1.9).

## Discussion

### Criteria ranking

In breeding programs, the mating design mainly relies on crosses involving the best parental inbred lines so as to ensure high mean progeny performance. The problem is that the highest-performing individuals may bear similar sets of alleles and actually produce fewer outstanding progenies (with less genetic variance) than the parents that have fewer yet favorable complementary alleles. As it is not feasible to evaluate all possible crosses in the field, it would be valuable to be able to predict the value of a cross or mating design in advance. Instead of focusing on the parental performance, the idea is to estimate a proxy of the value of top progenies based on the predicted progeny mean and variance. There have been several not very successful attempts made using distances between parents based on phenotypes (Souza and Sorrells 1991; Utz et al. 2001), genetic distance based on molecular markers (Bohn et al. 1999; Hung et al. 2012), molecular scores (summing QTL effects) and GEBV (summing marker effects estimated by ridge regression) (Tiede et al. 2015). The idea of genomic mating (GM) strategies is to use the genomic cross selection criterion (CSC) to optimize the complementation of parents to be mated (Akdemir and Isidro-Sánchez 2016). As progeny genetic variance is generated by randomly sampling recombinations between parental chromosomes and then sampling those chromosomes during meiotic division, if we could accurately estimate marker effects as well as recombination rates between markers, then it would be possible to optimize mating by maximizing the probability of obtaining individuals that cumulate a maximum of favorable alleles. Several CSC have been proposed to rank the crosses that are focused on different properties of the right-hand tail of the predicted distribution of progeny breeding values: UC (expected mean value of top progeny), PROBA (expected percentage of progeny with genetic value higher than a threshold), EMBV (best progeny value among N progenies) and OHV (best theoretical progeny value). One goal of the present study was to rank criteria based on their ability to provide superior genetic gain, but also to assess their impact on genetic diversity in order to measure their long-term genetic gain sustainability. Even with unperfect marker estimation, mating designs optimized using progeny variance estimates provided superior genetic gain and/or superior genetic diversity in progeny than mating designs solely optimized with regard to parental breeding values.

The PM criterion served as a reference. For all scenarios, other CSC (except OHV) provided superior genetic gain in top progeny when selection was stringent. Moreover, the PM criterion never offered the best trade-off between genetic gain and genetic diversity in the 7% top progenies when applying diversity constraints on parents (**Figures 2 ad 3**) or when not applying any (**Figure 4**). The set of best trade-offs was quite similar across scenarios. The OHV criterion systematically belonged to this set in the pareto curve and was associated with a minor genic variance loss but also the lowest genetic gain. The potential of OHV to maintain genetic diversity has already been demonstrated by Daetwyler et al. (2015). PROBA, UC3, UC1 and EMBV criteria provided the best trade-offs and were associated with the highest genetic gain, whereas there was a genic variance loss close to that observed using PM (up to 4% more genic variance).

This study tested the impact of two CSC, PROBA and UC3 adapted from recent literature. The PROBA criterion, as described by Wellmann (2019), ranked crosses based on their probability of producing progeny superior to the best parental line. PROBA provided among the highest elite progenies for all scenarios (**Figure 2**), except for TRUE + unselected and NO CONSTRAINT scenarios (**Figure 4**) where UC (criterion based on the expected superior quantile value of the progeny distribution) worked better. Note that a threshold must be set for the PROBA criterion. In this study, we opted to set this threshold according to the genetic value (TBV for TRUE scenarios or GEBV for ESTIMATED scenarios) of the best parental line. However, other authors have suggested using different thresholds (Wellmann 2019), e.g. the genetic gain to reach at next generation. Note that if the threshold is too high (or too low) compared to the expected progeny distributions (for a cross population), most crosses will have a 0 (or 1) PROBA value, which complicates optimization of this criterion.

The UC3 criterion aims to maximize the expected value of the 7% best progenies of the whole mating design. It is a direct application of the Index 5 criterion concept tested in Bijma et al. (2020) but with no numerical approximation, thereby increasing the computation time. In terms of genetic gain, the UC3 criterion was among the best CSC for NO CONSTRAINTS scenarios and TRUE + unselected scenarios with CONSTRAINTS. Note that for UC3 and EMBV, we could not use linear programming as for other CSC, so it was much more compute-intensive. For instance, it took < 10 min to optimize a mating design between 35k candidate crosses (pre-selection of the 10% crosses with the highest PM:), < 5 h to choose between 350k crosses (no pre-selection) using linear programming and around a day for UC3 or EMBV to reach reasonable convergence with our homemade genetic algorithm.

In conclusion, UC1 and UC2 criteria are a good trade-off for quick genetic gain optimization while maintaining genetic diversity.

### Factors influencing the added-value of CSC compared to PM

According to **Figures 2 and 4**, the relative increase in progeny TBV for CSC based on progeny variance estimation was significant for TRUE scenarios but not for ESTIMATED scenarios. For TRUE scenarios, CSC were more efficient for selected compared to unselected scenarios. Two non-exclusive factors could explain these results.

#### Progeny variance estimation accuracy

First, CSC using progeny standard deviation estimates (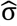) was hampered by higher estimation error than the conventional PM criterion based solely on progeny mean estimates 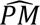 (**Table 2**). The correlation between estimated (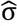) and true standard deviation (σ) was on average 4 to 22 points lower than the correlation between the estimated (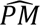) and true parental mean (PM). Note that both 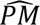 and 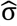 accuracies were higher in ESTIMATED + unselected populations than in ESTIMATED + selected populations. It is hard to determine if it is due to the lower heritability in selected populations (because the environmental variance was set as constant during *in-silico* breeding) or the negative correlation between QTLs (Bulmer effect).

**Table 2:** Correlation of the expected mean progeny estimate 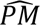 and progeny standard deviation 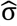 compared to their true PM and σ values. Marker effects were estimated using GBLUP for selected and unselected scenarios. Values were computed on the 10% crosses with the highest PM. Heritability was computed as the ratio between twice the genetic variance parameter estimated by GBLUP and the phenotypic variance.

| Population | $\widehat{h^2}$ | $\text{cor}(\widehat{PM}, PM)$ | $\text{cor}(\widehat{\sigma}, \sigma)$ |
| --- | --- | --- | --- |
| Unselected | $0.4 \pm 0.06$ | $0.45 \pm 0.05$ | $0.41 \pm 0.07$ |
| Selected | $0.3 \pm 0.03$ | $0.38 \pm 0.04$ | $0.16 \pm 0.07$ |

The lower accuracy of progeny variance estimates compared to genetic values has been reported in many studies (Neyhart and Smith 2019, Adeyemo and Bernardo 2019, Wolfe et al. 2021, Santos et al. 2019). Factors influencing the GEBV estimation accuracy, e.g. phenotyping quality, experimental design, statistical model used to take environmental effects into account, as well as the genetic relationship between the candidate and training population, probably impact progeny variance estimation accuracy as well. Concerning GEBV estimation, the different genomic selection models tested in the literature usually lead to slight or moderate improvement in GEBV accuracy for quantitative traits, while sometimes providing a significant improvement when trait variations were controlled by a few heterogenous QTLs (Daetwyler et al. 2008; Heslot et al. 2012). However, for progeny variance estimation, Bayesian models may markedly improve the accuracy for quantitative traits compared to the GBLUP model because of their ability to model the heterogeneity or uncertainty of marker effects. For example, Lehermeier et al. (2017) suggested using an Markov chain Monte Carlo (MCMC) algorithm to calculate the posterior mean of progeny variance (Sorensen et al. 2001, Lehermeier et al. 2017). In matrix notations, the progeny variance is calculated as 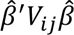, where 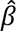 is the vector of estimated markers effects and *V_ij_* is the variance-covariance matrix of marker genotypes of the progeny derived from the cross between *Parent_i_* and *Parent_j_*. The MCMC algorithm allows estimation of the posterior distribution of 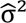 by averaging the product 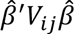 for each sample of the posterior distribution of 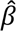 (posterior mean variance [PMV] estimates). Such PMV estimates were shown to be quite accurate in estimating the true progeny variance. For instance, in simulations run by Lehermeier et al. (2017), for h² = 0.4 with a 100–600 training population size range, the bias in the PVM estimate of progeny variance 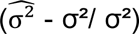 ranged from 0.06 to 0.21, while the correlation with the true value ranged from 0.58 to 0.65. This was much more accurate than what we obtained with our data for a similar scenario (unselected + ESTIMATED, h² = 0.4, training set size = 835, GBLUP model) with an average −0.82 ± 0.04 bias and 0.41 ± 0.07 correlation. Using empirical data, Wolfe et al. (2021) obtained a PMV estimate correlation with a real phenotypic observation of progeny variance ranging from 0 to 0.4 for different traits. Another strategy for estimating marker effects is to use selection models such as Bayesian Lasso that basically remove markers having very minor effects. In Santos et al. (2019) and Tiede et al. (2015), the Bayesian Lasso model provided more accurate marker effects and progeny variance estimates than GBLUP, but this was not the case in Yao et al. (2018). Finally, other GS models could be interesting to test with regard to increasing the progeny variance estimate accuracy, e.g. models using haplotypic blocks instead of markers (Cole and VanRaden, 2011, Bonk et al., 2016). The idea is that combinations of alleles in haplotypic blocks may be better estimated (if present in the training population) than individual SNPs and segregate as a block in progeny. For bread wheat, the recombination hotspots described in Danguy des Déserts et al. (2021) could be used as haplotype block separators, for instance.

#### Progeny variance variability of candidate crosses

The benefits of CSC based on progeny variance estimation also highly depend on the ratio between the progeny standard deviation and progeny mean variance t = var(σ)/var(PM). To understand why, let us follow the reasoning of Zong & Jannink (2007) based on an example with the UC criterion: the expected value of the superior fraction q of the progeny of a cross is computed as UC = PM + i*σ, with i being the selection intensity corresponding to the selected quantile q. The variance of UC values is thus equal to var(UC) = var(PM) + i²*var(σ) + 2*i*cov(PM, σ). We can thus hypothesize that the lower the t ratio, the more the UC variance could be explained by the PM variance. In other words, when the t ratio is low, the genetic values of parent (e.g. ∼ PM) values drive the expected superior progeny value. Hence, all CSC tend to select the same crosses, leading to a low additional genetic gain of alternative CSC over PM. For ESTIMATED scenarios, the t ratio was 4-fold lower than for TRUE scenarios (**Table 3**). As expected, the mating designs in our analysis converged when the t ratio decreased. **Supplementary Figure S2** shows a higher pairwise correlation of criteria for all candidate crosses for ESTIMATED (t = 2-3%) compared to TRUE (t = 6-11%) scenarios. This means that best parent ranking tended to be more similar for ESTIMATED compared to TRUE scenarios. Second, the proportion of shared parents between mating designs obtained with different criteria also increased for ESTIMATED scenarios, as well as the genetic similarity between recruited parents (**Supplementary Protocol S3, Supplementary Figures S1 and S3**). This means that a more similar cohort of parents was indeed selected across CSC. As the t ratio is highly decisive for the added value of alternative CSC over PM, it is important to properly estimate the progeny variance, e.g. using previously described models (PMV, Bayesian Lasso).

**Table 3:** Ratio between the variance of progeny standard deviation σ and the progeny expected mean PM for various scenarios. This was calculated for the 10% crosses with the highest PM.

| TRUE<br>$\text{var}(\sigma)/\text{var}(\text{PM})$ | | ESTIMATED<br>$\text{var}(\hat{\sigma})/\text{var}(\widehat{PM})$. | |
| --- | --- | --- | --- |
| Unselected | Selected | Unselected | Selected |
| 6% ± 1% | 11% ± 2% | 3% ± 0.1% | 2% ± 0.2% |

Bijma et al. (2020) tested the added value of several CSC for populations with different t ratios, and populations with a high t ratio systematically showed higher alternative CSC benefits. We can list some population types that are expected to have a high t ratio, and would thus be worthy of CSC implementation. First, Populations that experienced selection. In Bijma et al. (2020) it was hypothesized that in the context of an infinitesimal model and infinite populations the progeny variance does not change over generations. However, in our simulations, e.g. in finite populations with a finite number of causal loci, var(PM) was indeed reduced by selection (3-fold lower for selected scenarios compared to unselected scenarios), but var(σ) was also reduced (1.4-fold lower for selected compared to unselected scenarios). But the TRUE t ratio still increased, along with the expected benefits of CSC based on progeny variance estimation.

Second, structured populations can also lead to high t ratios. Structured populations arise when crossing elites with genetic resources (GR) from different genetic groups, in pre-breeding programs for instance, or to a lesser extent when crossing elite parents to elites from different breeding companies. When crossing parents from two highly differentiated populations, the t ratio may increase because of a higher magnitude of var(σ). Genetic differentiation leads to higher polymorphism between parents from different genetic groups (Wahlund effect) and among progenies, and thus higher progeny variance. According to Bijma et al. (2020), structuring in plants may explain the negative correlation between PM and σ reported in several publications in maize (Bernardo 2014; Mohammadi et al. 2015), bread wheat (Lado et al. 2017) and barley (Abed and Belzile 2019; Neyhart and Smith 2019). In our case, we also observed a negative relationship between PM and σ in INRAE-AO data analyses and simulations. The negative relationship was greater in unselected scenarios (**Supplementary Figure S2**). A negative correlation indicated that crosses with a low to medium parental mean (elite*GR) had a higher progeny variance than crosses between elite parents. In these situations, ranking crosses according to CSC based on progeny variance estimation may thus be very useful for increasing genetic gain.

We hypothesized that the genetic structure of the population and accuracy of progeny variance estimates were the two factors explaining high t ratios, and in turn high benefits of alternative CSC, in the plant breeding programs of Lehermeier et al. (2017) and Yao et al. 2018. Lehermeier et al. (2017) used a maize NAM population built with European dent landraces (Bauer et al. 2013) crossed to one elite accession, leading to a family-structured progeny. The ratio var(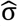)/var(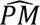) was on average 14% in the findings of Lehermeier et al. (2017) (Table 2 in their article, h² = 0.2 and h² = 0.6). The ratio obtained in our study using elite bread wheat material ranged from 2 to 11% (**Table 3**). In Lehermeier et al. (2017), the genetic gain provided by UC compared to PM was > 0.2 genetic standard deviation (σ_*g*_) at a < 10% selection rate (Figure 4 in their article). This was 5-fold higher than our best results for the 7% top progenies under similar scenarios (unselected + TRUE populations: genetic gain = 0.04 σ_*g*_; unselected + ESTIMATED, h²=0.4: 0.035 σ_*g*_). Yao et al. (2018) used bread wheat crosses involving Chinese and Australian lines that were likely very differentiated and thus likely associated with a high t ratio. In Yao et al. (2018), the genetic gain provided by UC was 0.06 σ_*g*_ at h² = 0.3, 0.08 σ_*g*_ at h² = 0.5 σ_*g*_ and 0.13 σ_*g*_ at h² = 0.8, for a selection rate ranging from 1 to 10%. This level was similar to what we observed for TRUE + unselected scenarios and 2-fold more compared to ESTIMATED + unselected scenarios (0.035 σ_*g*_).

### Trade-off between genetic gain and genetic diversity

The breeder’s equation implies that genetic gain is proportional to the selection intensity and genetic variance. However, the theory also predicts that in an isolated breeding program, without extrinsic germplasm introduction, each selection step is associated with a reduction in genetic variance. Genetic gain in successive generations would thus be expected to decrease, and finally converge to 0 when there is no longer genetic variance in the breeding population (Jannink et al., 2010). This phenomenon is faster with genomic selection, which decreases the generation interval, increases the selection intensity if the accuracy is high, and increases the probability of selecting related individuals (Clark et al. 2011, Pszczola et al. 2012).

Several methods have been suggested in the literature to come up with a trade-off between genetic gain and genetic diversity. For example, several authors (Jannink et al. 2010; Goddard 2009; Hayes et al. 2009) suggested giving more weight to rare and favorable alleles when computing GEBV on candidate parents (weighted genomic selection [WGS]). Alternatively, Goiffon et al. (2016) suggested selecting a set of candidate parents that bear at least one copy of all beneficial alleles. These two methods rely on marker effects estimated by genomic predictions. These estimates are expected to change over the generations due to selection, recombination and the resulting variations in LD between markers and QTLs. Alternatively, the optimal contribution selection (Meuwissen 1997) or optimal cross selection (Kinghorn et al. 2009; Allier al. 2019, OCS) methods are geared towards optimizing parental contributions to progeny in order to maximize genetic gain while constraining average pairwise inbreeding, which is proportional to the genetic variance loss (Falconer and Mackay 1996, reviewed in Woolliams et al. 2015). In plants, these methods have been adapted to inbreds by Allier et al. (2019) to maximize genetic gain while limiting the loss of mean expected heterozygosity in future progeny: 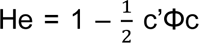 where c is the contribution of parents to progeny, Φ is the identity by state matrix (Allier et al. 2019). The expected genetic diversity He is determined by two factors, i.e. the distribution of progenies among candidate parents (c values) and the genetic similarity of parents Φ. As a rule of thumb, the overuse of the few (best) parents (Wray and Thompson, 1990) and the use of highly similar parents have a negative impact on the expected heterozygosity. In this study, instead of controlling He in progeny, we used more simple practical CONSTRAINTS on the mating design that forced us to allocate progenies to many crosses and parents (by imposing a minimum number of parents and crosses) and hampered crossing of similar parents. The CONSTRAINTS were likely less accurate in managing genetic diversity than OCS because they did not explicitly include the co-ancestry of parents. Yet they had the advantage of being easy to define as they could be based on usual breeding practices and budget constraints. In INRAE-AO material, we showed that setting constraints on parental contributions actually had little impact on genetic gain but highly preserved the genetic diversity.

## Conclusion

For an elite bread wheat breeding program, crossing parents with the highest genetic values is likely not the best means to maximize the usefulness of progeny. Alternative cross selection criteria that take progeny variance estimation into account could provide better elite progeny, improve the mean population value while maintaining more genetic diversity. However, the efficiency of these alternative CSC depends on the progeny variance estimation accuracy, which requires some improvement. The application of constraints to limit genetic diversity erosion when selecting crosses slightly decreased the genetic gain while preserving higher genetic diversity for long-term genetic gain. The use of such mating design optimization methods will be facilitated by the enhancement of genomic prediction methods and their widespread use in breeding programs. The implementation of such methods requires parental genotyping and a large phenotypic and genotypic database to predict marker effects, which is often carried out for genomic predictions of the *per se* value of advanced lines in breeding companies.

## Acknowledgments

The authors would like to thank the GenoToul bioinformatics platform Toulouse Occitanie (Bioinfo Genotoul, doi: 10.15454/1.5572369328961167E12) for providing support, computing and storage resources. The authors are also grateful to Hélène Rimbert for her help in identifying SNP positions on refSeq v1.0. The doctoral contract and activities of ADDD were funded by the INRAE metaprogram SELGEN and Florimond Desprez (Cappelle-en-Pévèle, France). Genotyping was supported by the Breedwheat grant (ANR-10-BTBR-03). The authors thank Simon de Givry and Daniel Ruiz for their advice on linear programming and genetic algorithms.

## Data Availability

Genotypes (<u>GenotypingData.txt</u>) and phenotypes are available in the INRAEDataverse repository (https://data.inra.fr/) with the following links https://doi.org/10.15454/AABGO7 and https://doi.org/10.57745/BSHZKV Scripts to reproduce all the results are available on Github (https://github.com/aldanguy/mating_plans_bread_wheat).

## List of acronyms

CSC: cross selection criteria
EMBV: expected maximum haploid breeding value
GA: genetic algorithm
GEBV: genomic estimated breeding value
GS: genomic selection
LD: linkage disequilibrium
OHV: optimal haploid value
PM: parental mean
PMV: posterior mean variance
PROBA: probability of a given cross progeny to exceed a given threshold (the best parental value here)
TBV: true breeding value
TP: training population
UC: usefulness criterion
UC1: expected mean value of a given cross top 7% progeny
UC2: expected mean value of a given cross top 0.01% progeny
UC3: expected mean value of a given cross progeny superior to the 93% quantile of the whole mating design

